# Maternal undernutrition exacerbates microbiota-driven growth stunting through pre and postnatal effects

**DOI:** 10.1101/2025.03.06.641866

**Authors:** Yadeliz Serrano Matos, Claire Williams, Jasmine Cano, Lindsey Bihuniak, Aria Kamal, Carrie A. Cowardin

**Affiliations:** Division of Pediatric Gastroenterology & Hepatology, Department of Pediatrics, University of Virginia School of Medicine, Charlottesville, Virginia 22908, USA; Department of Microbiology, Immunology and Cancer Biology, University of Virginia School of Medicine, Charlottesville, Virginia 22908, USA

**Keywords:** Stunting, microbiome, growth, undernutrition

## Abstract

Linear growth stunting due to undernutrition affects 20% of children under the age of five with far-reaching consequences, including increased susceptibility to infection and altered cognitive development. Current nutritional interventions are largely ineffective in rescuing linear growth. A significant proportion of stunting originates *in utero;* however, the mechanisms by which maternal undernutrition is transmitted between generations remain poorly characterized. Here we employed a gnotobiotic murine model of intergenerational undernutrition in which offspring are exposed to distinct microbial communities both *in utero* and after birth. We found that maternal undernutrition exacerbated offspring growth deficits in a microbiota-dependent manner. Maternal undernutrition also altered the offspring microbiome, increasing abundance of *Enterococcus* species and *Escherichia coli*. Cross-fostering demonstrated critical roles for the microbiota in growth during both gestation and early life. These findings emphasize the need to target both mother and child in the design of nutritional therapies for undernutrition.

## INTRODUCTION

Linear growth stunting (height-for-age Z score ≤ two standard deviations below the WHO median) is a manifestation of undernutrition that affects 20% of children under five years of age in low and middle-income countries (LMICs)^1^. Relative to wasting (characterized by low weight-for-age), stunting has far-reaching consequences throughout the lifespan, including impaired cognitive development, increased rates of infection, and an elevated risk of metabolic disorders^2^. Stunting is an intergenerational and multifactorial syndrome^3^, with stunted mothers at higher risk of giving birth to underweight and pre-term newborns that experience stunting soon after birth^4,5^. Longitudinal studies indicate that the onset of stunting often occurs within the first three months of life, with the reversal of stunting after this critical time period being less likely^1^. Current nutritional interventions have had little success in rescuing stunted growth^6^, indicating the importance of understanding additional factors beyond nutrition that lead to the development and persistence of stunting.

Environmental Enteric Dysfunction (EED) is pervasive in undernourished populations and may in part explain the poor effectiveness of current nutritional therapies^5^. EED is characterized by epithelial barrier dysfunction and malabsorption that is thought to be a consequence of microbiome disruption, pathogen carriage and accompanying intestinal inflammation^7^. The microbiome plays a crucial role during undernutrition^8–12^. However, the mechanisms by which microbial communities shape host growth are still being explored. Previous studies have demonstrated that undernourished Malawian children harbor microbiota that are chronologically immature compared to healthy controls^11^. Colonization of germ-free mice with these communities lead to impaired growth, altered bone morphology, and metabolic abnormalities^11^. These results suggest compositional changes in the microbiome of undernourished children are linked to long-term growth deficits. Because a significant proportion of childhood stunting emerges *in utero*^4^, an emergent hypothesis in the field suggests maternal microbes may also shape long-term growth and immunity during pregnancy and early life^3^. Maternal health is critical for healthy fetal development, and studies from high income countries suggest intestinal inflammation can lead to adverse birth outcomes. These findings have led to a renewed focus on maternal intestinal function during pregnancy in LMICs.

We previously demonstrated that intergenerational colonization with microbes from stunted infant donors leads to the development of stunting-like features in a gnotobiotic mouse model of undernutrition^13^. Our findings raised critical questions about when in life growth deficits arose and whether they could be rescued by exposure to healthy microbiota later on. To investigate these questions, we first explored the role of maternal undernutrition in offspring growth. We found that maternal exposure to a nutrient-deficient diet significantly impacted offspring survival in the context of a stunted donor (SD) but not a healthy donor (HD) microbiota. Maternal diet exposure also significantly impacted offspring linear and ponderal growth, intestinal physiology, and metabolism. Notably, maternal undernutrition significantly impacted the composition of the offspring microbiota, leading to increased abundance of *Enterococcus* and *Escherichia coli*. We next used two different cross-fostering approaches to determine the contribution of the maternal microbiota *in utero* and in the postnatal period. We observed that prenatal and postnatal exposures to SD microbiota both contribute to offspring development, with distinct effects on ponderal versus linear growth. This study sheds light onto the role of maternal nutrition on offspring growth and microbiota development, revealing the importance of targeting diet and microbiota-directed interventions throughout early life.

## RESULTS

### Maternal undernutrition exacerbates offspring growth stunting

To investigate the effects of maternal undernutrition on offspring growth, we modified our intergenerational model of undernutrition to include maternal diet and microbiota exposure in early life. In our original model, 4-week-old germ-free sires were colonized with microbiota derived from a healthy or a severely stunted Malawian infant donor^13^ and fed a protein and micronutrient-deficient diet (**Figure 1A**, **Figure S1A, Table S1**) representative of what is consumed by this donor population^11^. At 8 weeks of age, mice were switched to a nutrient-sufficient diet and 8-week-old germ-free females were introduced into the isolator and co-housed with sires as breeding pairs (**Figure S1A, Table S1)**. Offspring from these breeding mice were weaned into the same undernourished diet as the sires at 3 weeks of age. Because the female mice in this arm of the experiment were not exposed to the M8 diet, these offspring represent the HD-NMU (Healthy Donor, No Maternal Undernutrition) and SD-NMU (Stunted Donor, No Maternal Undernutrition) groups. To more closely model human undernutrition, we also included a second arm of the experiment in which both sires and dams were colonized with donor microbiota and exposed to the M8 diet from 4 to 8 weeks of age (**Figure 1B**). At 8 weeks of age, these animals were swapped back to the nutrient-sufficient diet and co-housed as breeding pairs. Offspring of these breeding pairs were weaned onto the M8 diet as described above. These animals represent the HD-MU (Healthy Donor, Maternal Undernutrition) and SD-MU (Stunted Donor, Maternal Undernutrition) groups. We hypothesized that direct colonization of dams and sires with donor microbiota and consumption of M8 diet for four weeks would exacerbate deficits in growth in both HD and SD groups.

**Figure 1.**
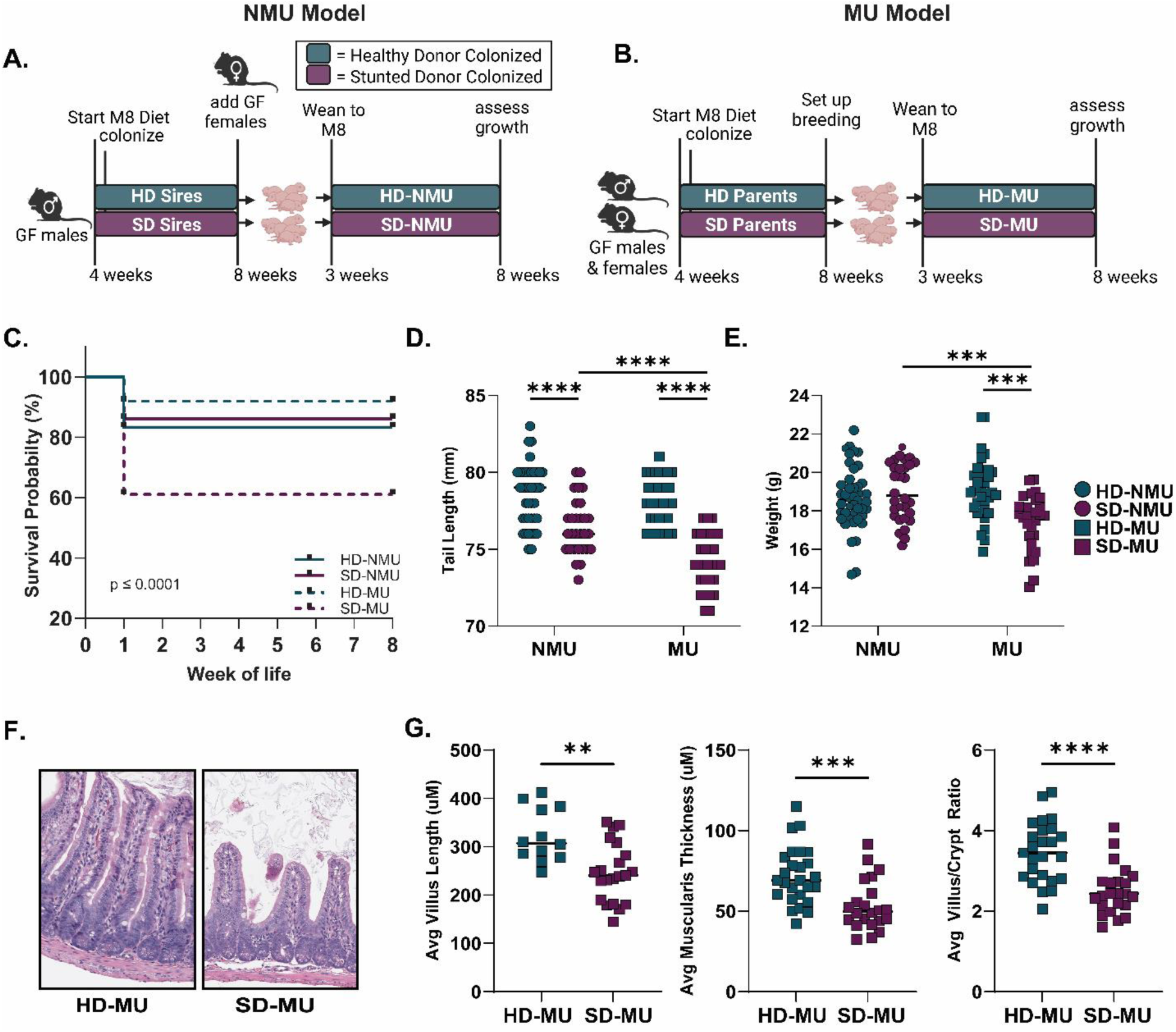
Maternal undernutrition exacerbates growth deficits in SD-MU mice. **(A-B)** Schematic of experimental design (created with Biorender.com). **(C)** Survival probability in both models. **(D)** Tail length at maturity. **(E)** Absolute body weight at maturity. **(F)** Representative histological images of H&E stained ileal tissue at 8 weeks of age. **(G)** Quantification of villus length, villus/crypt ratio and muscularis thickness in 8-week-old mice from NMU and MU models. Each data point represents an individual animal. * p ≤ 0.05, ** p ≤ 0.01, *** p ≤ 0.001, **** p ≤ 0.0001 by **(C)** Logrank (Mantel-Cox) test **(D-E)** Two-Way ANOVA with Holm-Šídák’s multiple comparisons test or **(G)** Mann-Whitney U test. **(C)** n=108-132 for NMU mice, n=136-149 for MU mice **(D)** n=29-40 for NMU mice, n=30-34 for MU mice **(G)** n = 12-21/group for Villus Length, n=22-25/group for Villus/Crypt Ratio and Muscularis Thickness.

Surprisingly, the introduction of maternal undernutrition into our model lead to a significant decrease in offspring survival in SD-MU compared to HD-NMU, SD-NMU and HD-MU pups (**Figure 1C**). This effect was reproducible across experiments (across four different iterations of the model – 8 dams/18 litters), with offspring mortality typically occurring within the first week of life. In addition to reducing offspring survival, maternal undernutrition also significantly exacerbated SD-MU growth stunting. At maturity, SD-MU offspring displayed shorter tail length compared to both HD groups as well as to SD-NMU offspring (**Figure 1D**). A similar trend was observed for overall weight at maturity (**Figure 1E**). These differences were not due to the number of pups per litter (**Figure S1B**) and were not sexually dimorphic (**Figure S1C-F**). These results suggest maternal undernutrition worsens growth deficits specifically in the presence of microbiota from a human donor with growth stunting, highlighting the important role of the early life microbiome in shaping long-term growth.

### SD microbiota consistently alters small intestinal morphology

In early life, small intestinal function is vital for growth by enabling nutrient absorption and physical and immune protection during the establishment of intestinal microbial communities^14^. Alteration in small intestinal tissue architecture is a hallmark feature of EED which is thought to contribute to malabsorption and perpetuate growth^15^. To investigate intestinal morphology, we performed histological scoring of H&E stained ileum tissue sections to assess established histological features of EED, including enterocyte injury and villous architecture^13,16^. We observed an increase in Enterocyte Injury and Villus Architecture scores in SD-MU offspring compared to HD-MU offspring at maturity (**Figure S2A**). We also assessed quantitative measures of small intestinal physiology, including villus length, muscularis thickness, and villus/crypt ratio. Results demonstrated a significant decrease in average villus length, villus/crypt ratio and muscularis thickness in the ileum of SD-MU compared to HD-MU offspring (**Figure 1F-G**). No significant difference in crypt length was observed between the groups (**Figure S2B**), suggesting the decrease in villus/crypt ratio in 8-week-old SD-MU mice was not due to crypt hyperplasia and instead was caused by a reduction in the length of the villi. None of the differences in intestinal morphology were sexually dimorphic (**Supplemental Figure 2C-E**).

Reductions in villus length have been associated with impaired gut function in stunted children and other mouse models of diet-dependent enteropathy^17–19^. Taken together, these results suggest that the SD microbiota drives reduced intestinal absorptive surface area in SD-MU offspring at maturity, potentially contributing to impaired growth^5^.

### Maternal undernutrition alters the abundance of specific taxa in the SD community

We next sought to investigate whether maternal undernutrition influenced the composition of the offspring microbiota at maturity. To address this question, we performed V3-V4 16s rRNA sequencing on the feces of offspring at 8 weeks of age. Principle Coordinate Analysis (PCoA) of Weighted UniFrac distances revealed that microbiota differences were largely driven by the donor microbiota received at birth (PERMANOVA, R^2^=.8742, p-value ** ≤ 0.001, **Figure 2A**, **Table S2A, S3A**). However, there was a significant difference in community composition due to maternal nutritional status within each donor context (p-value = 0.023, **Table S2A**). Interestingly, this result was primarily driven by microbiota composition in SD pups rather than HD pups (**Figure 2B**, **Table S2B-D**). We next calculated alpha diversity by Shannon Diversity Index. Although the SD community had higher diversity overall, there was no significant difference in diversity between SD-NMU and SD-MU offspring (**Figure 2C**). However, when we quantified the number of ASVs present in each group, we observed that both SD and HD MU offspring had significantly more ASVs detected compared to offspring of non-malnourished dams (**Figure 2D**).

**Figure 2.**
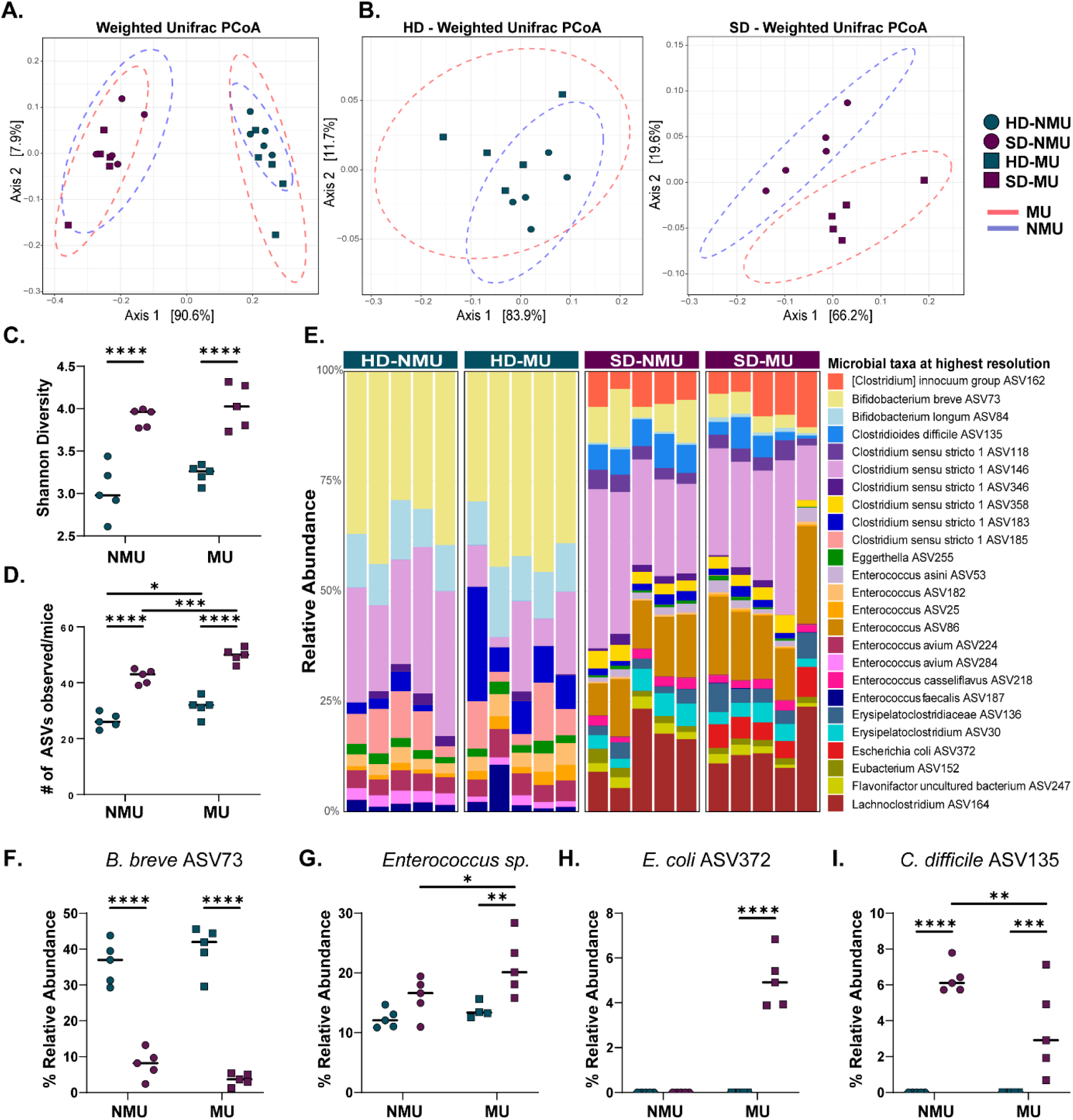
Maternal undernutrition alters the SD microbiota at maturity. Principal Component Analysis (PCoA) of Weighted Unifrac distances of the fecal microbiota measured by V3-V4 16s rRNA sequencing in **(A)** NMU and MU offspring of both donors **(B)** HD offspring and SD offspring individually. **(C)** Stacked bar plot of the 25 most abundant taxa across NMU and MU mice. **(D)** Shannon Diversity Index for NMU and MU mice. **(E)** Absolute number of observed ASVs in NMU and MU mice**. (F) %** Relative abundance of *Bifidobacterium breve,* **(G)** *Enterococcus spp.,* **(H)** *E. coli*, and **(I)** *Clostrodioides difficile* in NMU and MU animals. * p ≤ 0.05, ** p ≤ 0.01, *** p ≤ 0.001, **** p ≤ 0.0001 by **(A-B)** PERMANOVA and Pairwise Adonis or **(D-I)** Two-Way ANOVA with Šídák’s multiple comparisons test **(A-I)** n=5/group [3 females and 2 males].

We then explored differences in community composition in further detail (**Figure 2E**). Major compositional differences between the SD and HD groups included lower relative abundance of *Bifidobacterium breve* in the SD community (**Figure 2E**), which did not change significantly based on maternal nutritional status (**Figure 2F**). However, maternal undernutrition did significantly increase the relative abundance of *Enterococcus* species as well as *Escherichia coli* while reducing the abundance of *Clostridium difficile* in SD-MU relative to SD-NMU offspring (**Figure 2G-I**). Thus, maternal undernutrition significantly impacted specific taxa within the SD community without altering the HD community profile. In particular, maternal undernutrition increased the abundance of two potential pathobionts within the SD microbiota. Pathogenic *E. coli* are pervasive in undernourished populations and are capable of driving intestinal inflammation^20^ while *Enterococcus spp*. can become pathobionts in the immunocompromised host^21^. Although we cannot identify these ASVs as definitive pathogens based on these sequencing results, our findings highlight the importance of future investigation to determine what role *Enterococcus spp*. and *E. coli* play in shaping early life growth outcomes in this model.

### SD-MU offspring are born underweight and develop linear growth deficits after weaning

Because SD-MU offspring showed reduced survival soon after birth, we hypothesized that growth deficits would occur in the early postnatal period. To explore this possibility, we measured linear growth weekly starting in the first week of life. SD-MU offspring displayed significantly reduced tail length compared to HD-MU offspring starting at 4-weeks of age, one week after weaning onto the M8 diet (**Figure 3A**). In contrast, SD-MU pups had significantly decreased absolute body weight starting in the first week of life (**Figure 3B**). Rates of weight gain between SD-MU and HD-MU pups did not significantly differ over this time period (**Figure 3C**), suggesting that initial weight deficits in SD-MU pups were driving the observed reductions in weight at later time points. We next measured body weight in 1-day-old neonates, and again observed a reduction in body weight in SD-MU pups (**Figure S3A**). These results suggested that prenatal factors could be driving reduced ponderal growth in this group.

**Figure 3.**
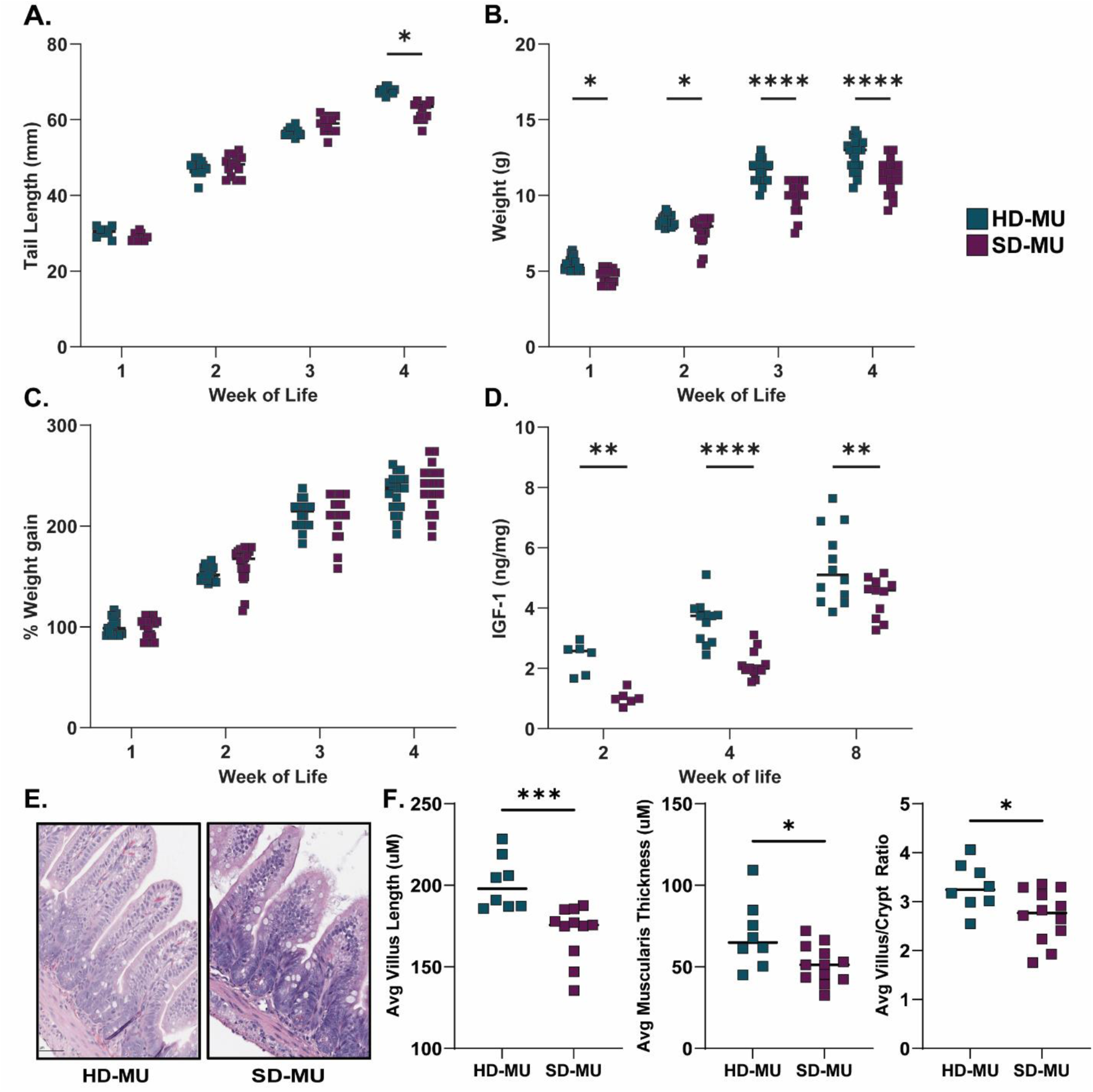
Microbiota composition in early life shapes physiological development. **(A)** Tail length and **(B)** absolute body weight in early life. **(C)** Percentage of weight gain over time. **(D)** Measurement of IGF-1 in liver tissue by ELISA in offspring from birth to maturity. **(E)** Representative histological images of H&E stained ileal tissue at 4 weeks of age. **(F)** Quantification of villus length, villus/crypt ratio and muscularis thickness in 4-week-old offspring from the MU group. Each point represents an individual mouse. Data shown includes equal number of female and male healthy donor colonized mice and stunted donor colonized mice unless otherwise specified. **(D)** 2-week-old SD-MU n= 4M/2F. **(F)** HD-MU n=9 (7M/2F), SD-MU n=12(5M/7F). * p ≤ 0.05, ** p ≤ 0.01, *** p ≤ 0.001, **** p ≤ 0.0001 by **(A-D)** Two-Way ANOVA with Holm-Šídák’s multiple comparisons test or **(F)** Mann-Whitney test.

Early life undernutrition is strongly correlated with reduced Insulin-like Growth Factor 1 (IGF-1) in humans, and IGF-1 levels are predictive of later growth^22,23^. This hormone is also influenced by intestinal microbes in other mouse models of undernutrition and growth stunting^24,25^. Because we observed major growth deficits in SD-MU offspring, we next measured IGF-1 in the liver of these mice. SD-MU offspring exhibited a significant reduction in liver IGF-1 protein levels at 2, 4 and 8 weeks of life (**Figure 3D**). Because linear growth deficits in SD-MU pups began at 4 weeks, we hypothesized SD-MU pups would exhibit features of EED at this time point. To explore this, we assessed histological features of the small intestine. SD-MU mice had a significantly higher histology score for both villus architecture and enterocyte injury compared to HD-MU mice at 4 weeks (**Figure S3B**). Quantification of villus length, villus/crypt ratio and muscularis thickness revealed a significant decrease for these three parameters in 4-week-old SD-MU relative to HD-MU mice (**Figure 3E-F**). These differences were not due to crypt hyperplasia and were not sexually dimorphic (**Figure S3C-F**). This suggests that as early as one-week post-weaning, SD-MU mice begin to develop intestinal dysfunction that may contribute to growth deficits over time.

### The weaning phase shapes community composition and the abundance of specific taxa

We next tracked microbiota establishment in early life to assess how these communities developed. We calculated weighted Unifrac distances (**Figure 4A**) for both HD-MU and SD-MU offspring, which showed that the age of the mice explained 92% of the differences within the HD group (**Table S2E**). Pairwise comparisons of microbiota composition overtime showed significant differences between all times points (**Table S2F**). Interestingly, differences in microbiota were explained the most by age (97%) when comparing HD-MU 2-week-old and 4-week-old mice (1 week before and after weaning), suggesting weaning and the introduction of the M8 diet strongly shapes the microbiota of HD-MU mice. In contrast, age explained a lower proportion of variance (64%) in SD-MU mice over time, although this result remained significant (**Figure 4B**, **Table S2G-H**).

**Figure 4.**
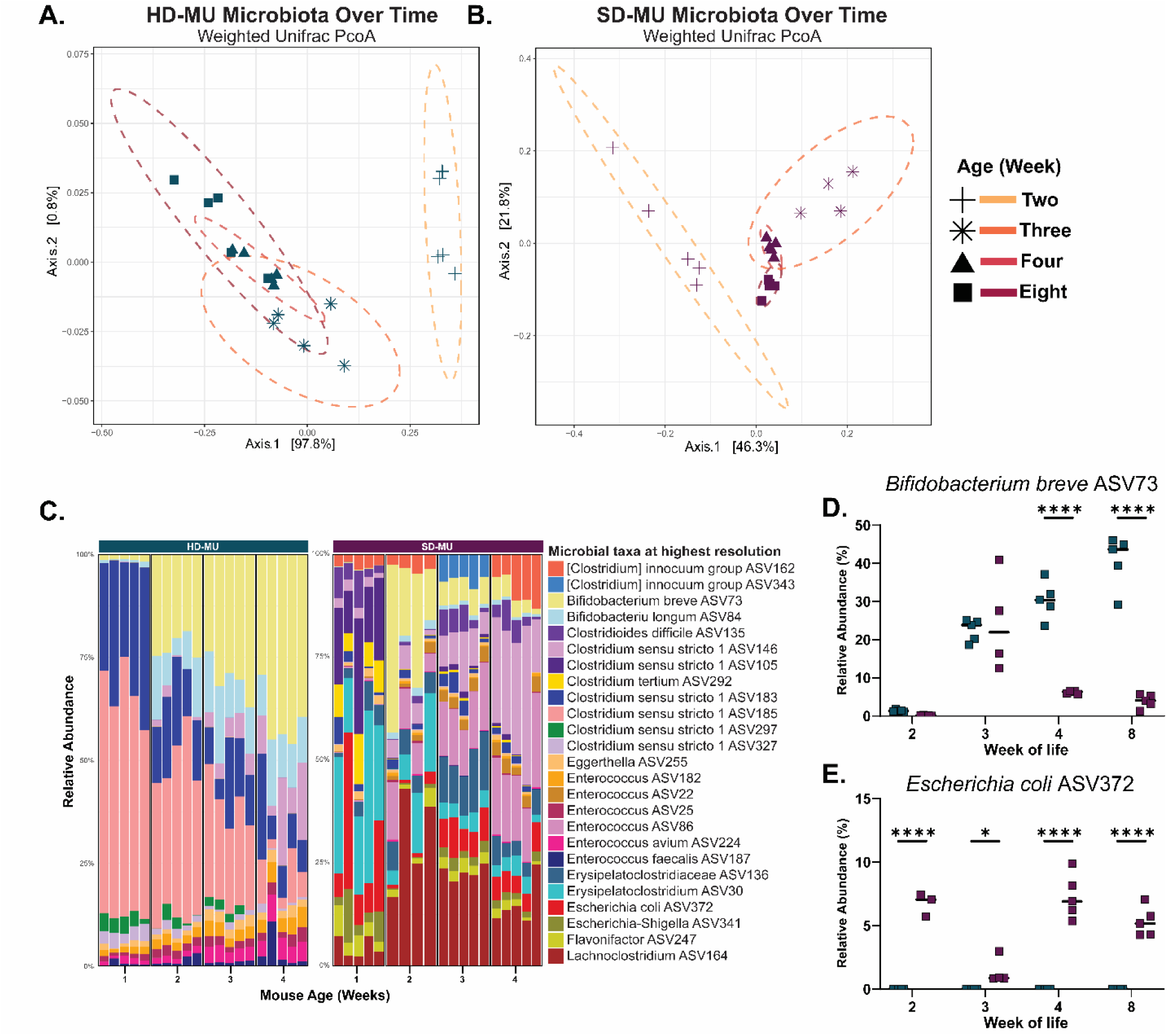
Development of the microbiota in early life. **(A-B)** Weighted Unifrac PCoA of the fecal microbiota measured by V3-V4 16s rRNA sequencing. **(C)** Stacked bar plot of the 25 most abundant taxa in HD-MU and SD-MU offspring in early life. **(D) %** Relative abundance of *Bifidobacterium breve* ASV73. **(E)** Relative abundance of *Escherichia coli* ASV372. Each individual point represent one mouse. * p ≤ 0.05, ** p ≤ 0.01, *** p ≤ 0.001, **** p ≤ 0.0001 **(A-B)** PERMANOVA and Pairwise Adonis **(D-E)** Two-Way ANOVA with Holm-Šídák’s multiple comparisons test. **(A-E)** n=5/group [3 females and 2 males].

We next explored differences in early life community composition in further detail. Major compositional differences between the SD-MU and HD-MU groups over time included higher absolute number of ASVs and increased alpha diversity in SD-MU offspring in early life (**Figure 4C**, **Figure S4A-B, Table S3B**). Interestingly, *Bifidobacterium breve* levels were comparable between 2- and 3-week-old HD-MU and SD-MU offspring. However, a significant decrease in this ASV was observed in SD-MU mice after weaning and the introduction of the M8 diet, while levels remained stable in HD-MU mice (**Figure 4D**). Furthermore, we observed a decrease in the relative abundance of *E. coli* in SD-MU offspring during weaning compared to pre-weaning and post-weaning time points (**Figure 4E, Figure S4C**). These results highlight the importance of the weaning transition in shaping the gut microbiota in this model.

### Postnatal microbiota exposure shapes linear growth

Recent findings suggest an undernourished microbiota can influence fetal development *in utero*^26^. Because we observed reductions in weight in SD-MU offspring beginning in the first week of life, we next sought to investigate the relative contribution of prenatal versus postnatal microbial colonization. To do so, we made use of a cross-fostering system in which germ-free offspring born to undernourished germ-free parents were fostered immediately after birth by either an HD-MU or SD-MU dam (**Figure 5A**). After cross-fostering, pups were once again weaned onto the M8 diet at three weeks of age. At maturity, germ-free mice cross-fostered to HD dams (GF-HD pups) displayed similar tail length compared to HD pups reared by HD dams. Similarly, germ-free mice cross fostered to SD dams (GF-SD pups) exhibited similar tail length to SD pups reared by SD dams (**Figure 5B**). Furthermore, GF-SD pups exhibited similar body weight to SD-MU pups at maturity, while GF-HD pups gain similar body weight compared to HD-MU pups (**Figure 5C**), suggesting postnatal exposure to the maternal microbiome or other maternal factors were sufficient to drive growth differences in these groups. In contrast to SD-MU animals, deficits in ponderal growth in GF-SD pups were observed starting at 1 week post-weaning (**Figure S5A**).

**Figure 5.**
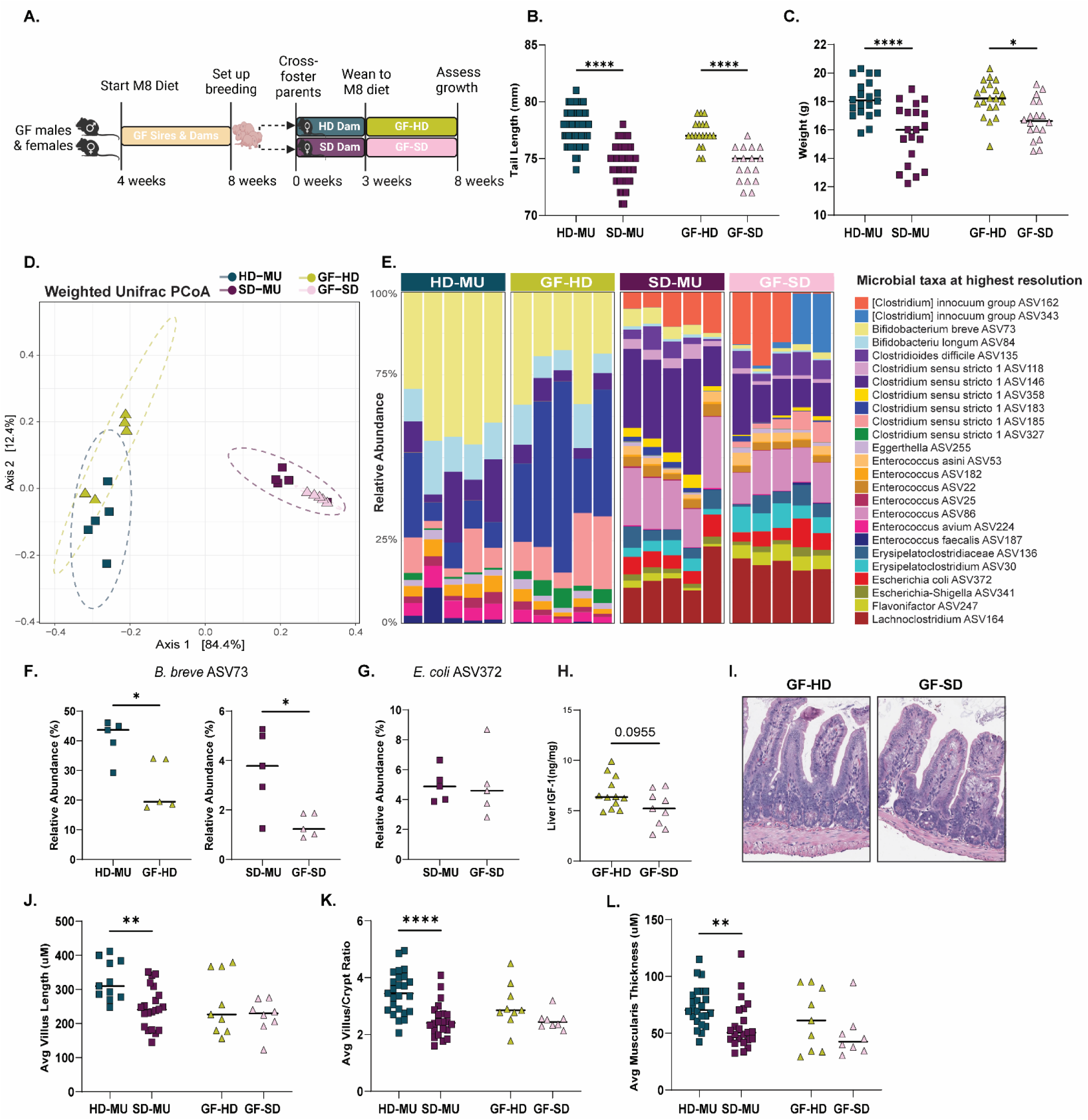
Postnatal exposure to SD microbiota is sufficient to drive long-term growth deficits. **(A)** Schematic of experimental design (created with Biorender.com). All measurements performed at maturity (8 weeks). **(B)** Tail length. **(C)** Absolute body weight. **(D)** Weighted Unifrac Principal Component Analysis (PCoA) of the fecal microbiome by V3-V4 16s rRNA sequencing. **(E)** 25 most abundant taxa across all groups. **(F)** Relative abundance of *Bifidobacterium breve* and **(G)** *Escherichia coli*. **(H)** IGF-1 levels by ELISA in liver tissue of GF-HD and GF-SD. **(I)** Representative histological images of H&E stained ileal tissue at 8 weeks of age. **(J)** Quantification of villus length. **(K)** Quantification of villus/crypt ratio. **(L)** Quantification of muscularis thickness. Each point represents an individual animal. **(J-K)** GF-HD n=9, 4F/5M, GF-SD n=9 4F/4M. MU mice n = 12-21/group for Villus Length, n=22-25/group for Villus/Crypt Ratio and Muscularis Thickness. * p ≤ 0.05, ** p ≤ 0.01, *** p ≤ 0.001, **** p ≤ 0.0001 by **(B-C, J-L)** Two-Way ANOVA with Šídák’s multiple comparisons test or **(F-H)** Mann-Whitney U test.

### Cross-fostered offspring show reduced abundance of Bifidobacterium breve

Due to the similarities in growth in GF-SD and SD-MU mice, we hypothesized that these animals would have a similar microbiome composition. Weighted UniFrac analysis of V3-V4 16S sequencing of the fecal microbiota at maturity demonstrated that cross-fostered GF pups clustered by donor with offspring born to colonized dams (**Figure 5D, E**), suggesting overall similar microbiome composition. However, pairwise comparisons demonstrated significant differences between HD-MU, SD-MU, GF-HD and GF-SD animals (**Table S2I-J**). Interestingly, when comparing the colonization of *Bifidobacterium breve* in cross-foster groups, we observed that both GF-HD and GF-SD mice exhibited reduced relative abundance of this ASV at maturity compared to HD-MU and SD-MU offspring (**Figure 5F. Table S3C**). In contrast, *E. coli* was present in GF-SD at a similar abundance to SD-MU mice (**Figure 5G**). These data could suggest that *Bifidobacterium breve* colonization may be more strongly shaped either by mode of acquisition (either at birth or in the postnatal period) or by prenatal maternal influences.

### Postnatal exposure is insufficient to shape some aspects of intestinal morphology

We next measured IGF-1 levels in cross-fostered animals and observed a trend towards a reduction of this protein in the liver of GF-SD mice compared to GF-HD mice at maturity (**Figure 5H**).To explore this further, we assessed the role of the postnatal microbiome in intestinal morphology by histological scoring of H&E stained ileum sections from cross-fostered GF offspring. Scores for enterocyte injury and villus architecture were increased in GF-SD compared to GF-HD mice (**Figure S5B**). Quantitative measurements of villus length and muscularis thickness at maturity revealed no significant differences in SD-MU, GF-HD, and GF-SD groups (**Figure 5I-L**). Similarly, we also did not observe a significant difference in average villus/crypt ratio between GF-HD and GF-SD pups (**Figure 5K**). We next measured average crypt length to determine if the decrease in villus/crypt ratio is a result of crypt hyperplasia, but did not observe significant differences (**Figure S5C**). Likewise, we did not observe differences in histological parameters by sex (**Figure S5C-F**). The results from these cross-fostering experiments suggest that exposure to microbes in the SD community immediately after birth is sufficient to drive reductions in linear and ponderal growth that manifest after weaning. In addition, germ-free cross-fostered mice show distinct intestinal morphology compared to SD-MU and HD-MU controls, suggesting prenatal exposure to the microbiota may contribute to these outcomes.

### Long-term growth outcomes are shaped by prenatal and postnatal exposures

To further explore the relative contribution of prenatal versus postnatal microbiota exposure in this model of undernutrition, we next employed a second cross-foster approach in which litters born to HD and SD dams were cross-fostered to the opposite dam at birth (**Figure 6A**). Consistent with previous experiments, pups were weaned onto the M8 diet at 3 weeks of age. At 8 weeks of age, these animals were euthanized to assess growth and changes in intestinal morphology. Tail length measurements at maturity revealed that SD, SD-HD (SD pups reared by HD dams) and HD-SD (HD pups reared by SD dams) pups exhibited significantly shorter tails compared to HD pups; however, HD-SD pups had significantly longer tails than SD-HD pups (**Figure 6B**). Interestingly, HD-MU and HD-SD pups had significantly increased absolute body weight at maturity compared to SD-MU and SD-HD pups (**Figure 6C**) while IGF-1 protein levels in the liver were significantly reduced in SD, SD-HD and HD-SD groups relative to HD pups alone (**Figure 6D**). These results suggest any exposure to the SD microbiota negatively impacted growth, but pups born to HD dams had slightly better growth outcomes than those born to SD dams.

**Figure 6.**
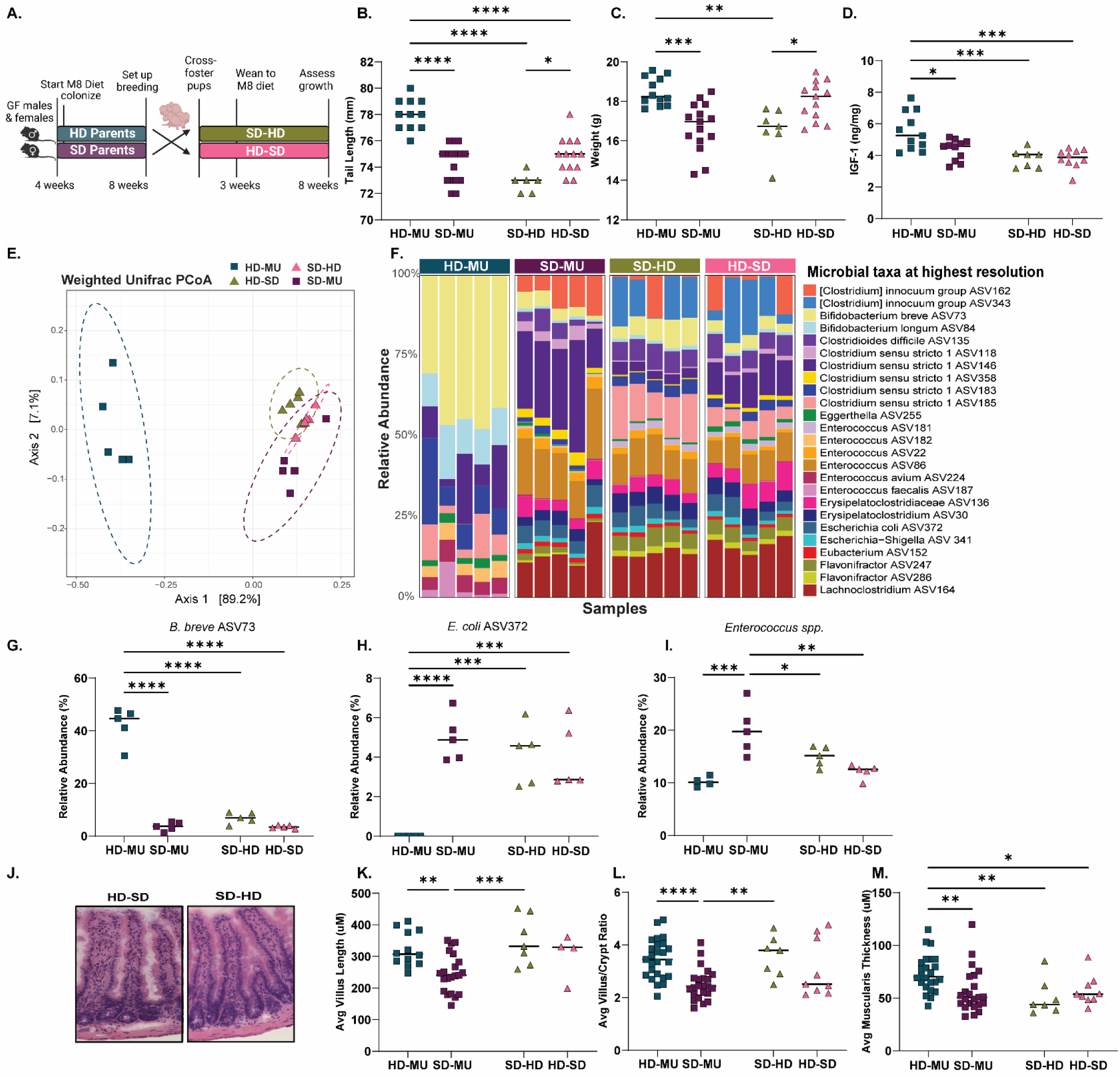
Postnatal exposure to SD microbiota negatively impacts linear but not ponderal growth. **(A)** Schematic of experimental design (created with Biorender.com). All measurements performed at maturity (8 weeks of age). **(B)** Tail length. **(C)** Absolute body weight. **(D)** IGF-1 by ELISA in liver tissue. **(E)** Weighted Unifrac Principal Component Analysis of the fecal microbiota measured by V3-V4 16s rRNA sequencing. **(F)** Stacked bar plot of the 25 most abundant taxa across groups**. (G)** Representative histological images of H&E stained ileal tissue. **(H)** Quantification of villus length. **(I)** Quantification of villus/crypt ratio. **(J)** Quantification of muscularis thickness. **(K)** Relative abundance of *Bifidobacterium breve* ASV43. **(L)** Relative abundance of *Enterococcus* spp. **(M).** Relative abundance of *Escherichia coli* ASV372. Each point represents an individual animal. * p ≤ 0.05, ** p ≤ 0.01, *** p ≤ 0.001, **** p ≤ 0.0001 by **(B-D, H-N)** Two-Way ANOVA with Šídák’s multiple comparisons test or **(E)** PERMANOVA and Pairwise Adonis comparisons.

We next investigated how this cross-fostering approach shaped the offspring microbiota. Analysis of Weighted UniFrac distances revealed HD-SD, SD-HD and SD-MU pups cluster closely together and form a distinct group compared to HD pups (**Figure 6E**), suggesting these animals have a more similar microbiome composition. However, we did identify significant differences in composition between all groups at maturity (**Table S2K**). Birth dam significantly influenced community composition (R^2^=0.25, p-value=0.001, **Table S2L**) as did the foster dam, with postnatal factors explaining slightly more of the differences at maturity (R^2^=0.34, p-value=0.001). Interestingly, HD-SD and SD-HD pups had a significant increase in the number of ASVs in the community, similar to SD-MU mice (**Figure S6A, Table S3D**). Furthermore, cross-fostered mice also had an increase in alpha diversity similar to SD-MU offspring (**Figure S6B**).

We also observed a reduction in the relative abundance of *B. breve* in HD-SD and SD-HD mice at maturity relative to HD-MU offspring, with a corresponding increase in *E. coli* (**Figure 6F-H**). Interestingly, cross-fostered offspring showed a significant decrease in the relative abundance of *Enterococcus* spp., suggesting exposure to the HD microbiota can reduce the abundance of these taxa (**Figure 6I**). Overall, these results suggest that the SD community largely overtakes the HD community in pups born to HD dams but reared by SD dams. This exposure significantly impacts offspring growth, resulting in HD-SD pups that have reduced tail length and IGF-1 relative to HD pups alone.

### Pre and postnatal factors influence intestinal morphology at maturity

As a potential underlying mediator for changes in growth, we next investigated whether different windows of microbiota exposure influenced intestinal morphology in cross-fostered pups. Histological scoring did not reveal significant differences in enterocyte and villus architecture for cross-fostered mice, although HD-SD mice did show reduced crypt length relative to SD-HD animals (**Figure S6C-D**). Interestingly, HD, HD-SD and SD-HD pups exhibited similar villus length that was significantly greater than that of SD pups, and villus/crypt ratio followed a similar trend (**Figure 6J-L**). However, we did observe a significant decrease in muscularis thickness for SD, SD-HD, and HD-SD pups compared to HD pups (**Figure 6M**). Thus, measurements of muscularis thickness in the small intestine appear to more closely correlate with growth outcomes in these animals. Overall, these results suggest microbiota exposure at distinct time points influences specific aspects of intestinal morphology.

## DISCUSSION

Our group previously developed a model of intergenerational undernutrition in which paternal and post-weaning undernutrition led to stunting-like features in offspring colonized with microbiota from stunted human infant donors^13^. While our previous work stresses the importance of understanding intergenerational influences on early-life growth, it did not explore a major hypothesized cause of its persistence — maternal undernutrition. Maternal nutritional status plays a key role in fetal growth restriction, low birth weights and poor postnatal growth, all of which are predictors of persistent growth stunting^27,28^. Furthermore, studies have shown that maternal undernutrition can lead to fetal epigenetic changes that influence metabolism, immune function and the microbiota in early life^29^. Therefore, investigation of the role of maternal undernutrition is critical to understanding the intergenerational transmission of growth stunting.

Indeed, the dominant line of thinking regarding nutritional therapies has pivoted towards maternal treatment in recent years^30^. However, major questions remain as to the precise timing and composition of these therapies. Because the impacts of undernutrition are both urgent and dire, the use of preclinical models to guide these approaches is critical. Here, we report that the introduction of maternal undernutrition exacerbated features of growth stunting and resulted in reduced linear and ponderal growth in offspring born to dams harboring microbiota from a stunted infant^13^. Interestingly, deficits in weight in SD pups were apparent from birth, while linear growth deficits developed post-weaning. These results mirror the development of stunting in children, who are frequently born underweight and go on to develop stunting within the first three months of life^1^. These observations demonstrate the utility of this model as a tool for the mechanistic study of how the early life diet and microbiota shape long-term developmental outcomes.

Maternal high fat diet exposure has been previously shown to shape the offspring microbiome and alter cognitive development^31,32^; however, the effects of maternal undernutrition on the offspring microbiome is less well understood. Our results demonstrate that maternal undernutrition shapes the abundance of specific microbial taxa at maturity, revealing persistent effects of maternal nutritional deficits. Taxa increased by maternal undernutrition included *Enterococcus* species as well as *E. coli*. Undernourished children often exhibit increased burden of *Escherichia* spp., with the presence of virulence genes in this taxon negatively correlated to height and IGF-1^33,34^. These findings highlight the importance of future studies to investigate microbial functions in further detail via more targeted and detailed analyses.

Recent work has also implicated the maternal microbiome in shaping offspring gastrointestinal development via expansion of intestinal stem cells, goblet cells, and enteroendocrine cells. This effect was not recapitulated in animals fostered by colonized dams but was instead dependent on the prenatal maternal microbiome^35^. Our findings support this conclusion and add further nuance on the role of maternal diet. In our model, intestinal morphology was shaped to a greater extent by prenatal factors, particularly with respect to muscularis thickness. Further understanding of signals regulating these changes could help identify additional pathways that regulate small intestine development to reveal targeted therapies that improve child intestinal absorptive function in early life.

Lastly, our cross-fostering studies showed that postnatal exposure to the stunted donor microbiota in germ-free pups was sufficient to recapitulate growth deficits observed in animals born to colonized dams. Results from a second cross-foster model using pups born to colonized dams suggested that any exposure to the SD microbial community negatively impacted growth, but offspring born to dams with healthy donor microbiota showed better growth outcomes. Both groups of cross-fostered offspring harbored microbiota that largely resembled the SD community, suggesting the SD microbiota was able to largely displace HD microbes acquired at birth. These results have major implications for the use of nutritional therapies in undernourished children and suggest that both pre and postnatal interventions may be required to durably restore growth in these populations.

### Limitations of this study

Due to the complexity and length of the experimental design employed, we were unable to investigate more than two microbiota donors. Future studies will be necessary to determine the generalizability of these results in additional human donor contexts. One potential mechanism to facilitate this goal involves the creation of a model community of microbes designed to capture common microbiota members present in many individuals within a specific population. Similarly, we were unable to full isolate the role of the prenatal maternal microbiome through the delivery of offspring by Cesarean section, allowing sterile neonates to be fostered by SD, HD or GF dams. This approach would more clearly delineate the role of the microbiota at each time point. While technically challenging to perform in gnotobiotic animals, this study would be extremely valuable and warrants further consideration.

## CONCLUSIONS

Maternal and child undernutrition continue to be major global health challenges. Children who are born stunted frequently remain so despite available nutritional therapies. Because linear growth is strongly linked to later health outcomes, investigating the origins of linear growth deficits in early life is critical to improving the effectiveness of current therapies and to developing new treatment approaches. Our model of maternal undernutrition allows for the mechanistic dissection of different developmental outcomes linked to the gut microbiota that are also observed in stunted children. These results have major implications for the use of nutritional therapies in undernourished children and suggest that both pre and postnatal interventions may be required to durably restore growth in these populations. In addition, this model provides a useful preclinical approach to test nutritional and microbiota-directed interventions at distinct life stages to optimize therapeutic benefit in low-resource settings.

### Resource availability

Demultiplexed V4 16 s rRNA sequencing reads have been deposited in the European Nucleotide Archive under accession number PRJEB86068. Sample metadata is provided in Supplementary Table S3<u>-B</u>. This manuscript does not report original code. All other data needed to evaluate the conclusions in the manuscript are available within the main text or supplementary materials.

## Acknowledgments

We thank the participants and investigators of the iLiNS-Dyad-M study for the samples used in this study. We also thank UVA’s Gnotobiotic and Germ-Free Animal Facility and Rory Laman for their help in performing these experiments, which would not have been possible without their dedicated efforts. We thank UVA’s Flow Cytometry Core Facility (RRID: SCR_017829) and Research Histology Core for their contributions to this work.

## Author contributions

Conceptualization, Y.S.M and C.A.C.; Sample collection, Y.S.M, J.C., C.W., L. B., A.K. and C.A.C.; Investigation, Y.S.M, and C.A.C., Analysis; Y.S.M, and C.A.C., Visualization, Y.S.M, and C.A.C.; Resources, Y.S.M, C.W., and C.A.C; Funding, Y.S.M. and C.A.C, Supervision, C.A.C; Writing, Y.S.M, and C.A.C.

## Declaration of generative AI and AI-assisted technologies

I hereby declare that no generative AI tools were used in the creation, analysis, or interpretation of this work. All data analysis and conclusions are the result of my own research and efforts, without the assistance of artificial intelligence systems.

## METHODS

### Donor microbiota and study protocol

Details of enrollment for the iLiNS-DYAD-M study were described in an earlier publication. Briefly, enrollment was open to consenting women over the age of 15 years with ultrasound confirmation of pregnancy of <20 weeks gestation in the Mangochi District of southern Malawi. The randomized controlled clinical trial [clinicaltrials.gov #NCT01239693] tested the effects of providing small quantity Lipid-based Nutrient Supplements (SQ-LNS) to pregnant and lactating women through 6 months postpartum and to their children through 6-18 months of age^36^. During pregnancy and 6 months thereafter, women received one daily capsule of iron-folic acid supplement (IFA group), one capsule containing 18 micronutrients (MMN group), or one 20-g sachet of SQ-LNS [lipid-based nutrient supplements (LNS), containing 21 micronutrients, protein, carbohydrates, essential fatty acids, and 118 kcal]. Children in the IFA and MMN groups received no supplementation; children in the LNS group received SQ-LNSs from 6 to 18 months^31^. Donors used in this study were selected from samples collected at the six-month time point (prior to supplementation) from a broader subset of donors based on their ability to colonize recipient gnotobiotic mice at an efficiency of >50% (more than half of taxa present in the original donor sample were identified in initial experiments). No significant differences in infant microbiota composition based on maternal supplementation at this time point were identified^37,38^.

### Gnotobiotic Mice

All gnotobiotic mouse experiments were performed using protocols approved by the University of Virginia Institutional Animal Care and Use Committee. All gnotobiotic animals used in this publication were germ-free C57BL/6NTac mice obtained from Taconic Biosciences. Upon arrival, germ-free status was verified by quantitative PCR. Mice were housed in plastic flexible film gnotobiotic isolators (Class Biologically Clean Ltd.) under a 12-hour light cycle. Animals received *ad libitum* access to food and water throughout the experiment, and were euthanized at the conclusion of the experiment using AVMA approved procedures. For experiments featuring maternal undernutrition, male and female mice were obtained from Taconic Biosciences at 4 weeks of age and transitioned to the M8 diet. Three days later, they were colonized with donor microbiota and maintained on M8 diet until 8 weeks of age. Males and females were co-housed and switched to an autoclaved nutrient-sufficient breeder chow (LabDiet 5021 Autoclavable Mouse Breeder Diet, LabDiet Inc.). For experiments without maternal undernutrition, male mice were obtained from Taconic Biosciences at 4 weeks of age and transitioned to the M8 diet. Three days later, they were colonized with donor microbiota and maintained on M8 diet until 8 weeks of age. At this point, 8-week-old germ-free females (also from Taconic Biosciences) were introduced into the donor isolators. Males and females were co-housed and switched to the autoclaved nutrient-sufficient breeder chow listed above.

For cross-fostering experiments, male and female mice were obtained from Taconic Biosciences at 4 weeks of age and transitioned to the M8 diet. Three days later, germ-free mice were split into three groups, each in a unique gnotobiotic isolator (HD, SD and germ-free). HD and SD mice were colonized with donor microbiota, while germ-free mice remained uncolonized. All three groups were maintained thereafter on the M8 diet until 8 weeks of age. At this time, they were all switched to breeder diet. Estrous cycles were synchronized by swapping male and female bedding. For germ-free cross-fostering experiments, once litters were born to both germ-free and colonized dams, neonates were transferred to their cross-foster HD or SD dam within three days. Neonates born to HD or SD dams at this time point were euthanized to prevent overcrowding.

Breeding mice were refreshed after six months, and offspring used in these experiments were derived from breeders from five separate rounds of colonization (total of 2-4 males and 6-8 females per colonization). Weight and tail length measurements were collected from birth to maturity. Tail length measurements in living mice were assessed using calipers sterilized to be used inside the isolators. The tail lengths measurements after euthanasia were performed using a standard laboratory ruler. All tail length measurements were done from the base of the tail to the tip along a straight line. For weight data collected in early life, animals were weighed weekly from 1 to 4 weeks of age inside the isolator. Weights taken inside the isolator were measured using a manual scale, while weights taken after euthanasia were measured on a digital scale outside the isolator. Neonatal weights were measured within 3 hours of birth for all mice inside the isolator using a manual scale. Fecal samples were collected at the indicated time points and immediately frozen. Samples were stored at -80°C until use. Data shown includes equal number of female and male healthy donor colonized mice and stunted donor colonized mice unless other wise specificied.

### Colonization and Diets

To prepare infant fecal samples for colonization of germ-free mice, aliquots of each sample were removed from storage at -80°C, weighed and immediately transferred into anaerobic conditions (atmosphere of 75% N2, 20% CO2 and 5% H2; vinyl anaerobic chambers from Coy Laboratory Products). Samples were subsequently resuspended in pre-reduced PBS containing 0.05% L-Cysteine Hydrochloride at a concentration of 10 mg/mL. Samples were vortexed for one minute and allowed to clarify by gravity for 5 minutes. The supernatant was removed to a fresh anaerobic tube and combined with an equal volume of sterile, pre-reduced PBS containing 0.05% L-Cysteine HCL and 30% glycerol. Gavage mixtures were aliquoted into sterile 2 mL screw cap tubes (Axygen) and frozen at -80°C until use. To colonize recipient mice, pools of gavage mixtures were sterilized externally with ionized hydrogen peroxide (STERAMIST System, TOMI Inc.) and passed into each isolator after appropriate exposure time (20 minutes). Animals were colonized via a single oral gavage with a 200 µl volume of gavage mixture.

The low protein and micro-nutrient deficient Malawi-8 diet was prepared as previously described^11,39^ and obtained from Dyets, Inc. Briefly, ingredients (corn flour, mustard greens, onions, tomatoes, ground peanuts, red kidney beans, canned pumpkin and peeled bananas) were cooked and combined in an industrial mixer. Dry pellets of the M8 diet were extruded, vacuum-sealed and double bagged prior to sterilization by irradiation (Steris Co). The nutritional content of the cooked and irradiated diet was assessed by N.P. Analytical Laboratories as described in Blanton et al^24^. LabDiet 5021 was sterilized by autoclaving at 129°C and 13.2 PSI for 15 minutes. Sterility of both diets was routinely assessed by culturing pellets in Brain Heart Infusion (BHI) broth (Millipore), Nutrient broth (Millipore), and Sabouraud-Dextrose (Millipore) broth for five days at 37 °C under aerobic conditions, and in BHI broth and Thioglycolate broth (Difco) supplemented with 0.05% L-Cysteine Hydrochloride (Sigma) under anaerobic conditions. After the five-day liquid culture, cultures of all diets were plated on BHI agar supplemented with sheep blood (Thermo Scientific). All diets were stored at -20 °C prior to use.

### Histopathology and Anthropometry

At the time of euthanasia, a 1 cm section of the proximal ileum was dissected from each mouse and fixed in 10% neutral-buffered formalin overnight at room temperature before being transferred to 70% Ethanol. Tissue processing and H&E staining were performed by the University of Virginia’s Research Histology Core or at Histowiz Inc. Samples were paraffin embedded and sectioned before mounting. Slides were stained with hematoxylin & eosin prior to imaging at a 20x magnification. To assess the histopathological features of the ileum, the stained tissues were scored in a blinded manner. Scores were assigned using a scoring system based off published findings in human intestinal biopsies ^35^. Scoring parameters consisted of 2 qualitative features: villous architecture (0, majority of villi are >3 crypt lengths long; 1, majority of villi are <3 but >1 crypt length long, with abnormality; 2, majority of villi are absent or <1 crypt length long, with abnormality) and enterocyte injury (0, majority of enterocytes show tall columnar morphology; 1, < 50% of enterocytes show low columnar, cuboidal, or flat morphology; 2, > 50% of enterocytes show low columnar, cuboidal, or flat morphology). Cumulative scores were calculated as the sum of the averaged score for both parameters. ImageJ was used to obtain three quantitative parameters consisting of ileum villus height (μm), ileum muscularis thickness (μm) and ileum crypt length (μm). Two measurements were obtained for villus length and crypts, three measurements for muscularis thickness parameter and averaged.

### Microbial Sequencing

Samples were shipped to SeqCenter for DNA extraction and sequencing. All standard DNA extractions at SeqCenter follow the ZymoBIOMICS™ DNA Miniprep Kit5. DNA was extracted from fecal samples following the guidelines in Appendix B of the ZymoBIOMICS™ DNA Miniprep Kit. Eluted DNA was then purified by running the effluent through the prepared Zymo-Spin™ III-HRC Filter. Final DNA concentrations were determined via Qubit. After DNA extractions, library prep was performed using Zymo Research’s Quick-16S kit with passed primers targeting the V3-V4 region of the 16S gene. The following sequencing primers were used: “CCTACGGGDGGCWGCAG” (341F) and “GACTACNVGGGTMTCTAATCC” (806R). After cleanup and normalization, samples were sequenced using a P1 or P2 600-cycle NextSeq2000 Flowcell to generate 2×301 bp paired-end (PE) reads. Quality control and adapter trimming were conducted with bcl-convert1 (v4.2.4). Primer-dimer sequences, identified as PCR artifacts, were removed from the FASTQ files using the following criteria: read length greater than 150 bp, PolyN strings with fewer than 10 consecutive Ns, and PolyG strings with fewer than 150 consecutive Gs

### Sequencing Quality Control & Data Analysis

Sequences were then denoised using Qiime2’s dada2 plugin3. Denoised sequences were assigned a microbial taxonomy by mapping to the Silva 138 99% ASVs full-length sequence database and the VSEARCH4 utility within Qiime2’s feature-classifier plugin. Prevalence filtering threshold was set to remove any ASVS ≤ 1.21% prevalence across all samples excluded from analysis. ASVs with a relative abundance across all samples ≤ 0.1% were removed as spurious ASVs. Phylogenetic trees for distance matrices were constructed using the following R packages: msa^40^ (version 1.38.0) and phagorn^41^ (version 2.12.1). Percentage of relative abundance were calculated by subsetting samples according to analyses required.

Ordination plots showing weighted UniFrac distances (a measurement of β diversity based on phylogenetic relatedness) were produced using the phyloseq package^42^ (version 1.50.0) in R. The stackbar plots shown in Fig. 2, Fig.4, Fig. 5 and Fig. 6 were created using the ggplot2^43^ in R (version 3.5.1).

### Protein Quantification

Liver tissue were collected at the time of euthanasia and flash frozen in liquid nitrogen. Liver samples were homogenized in Lysing Matrix F (MP Biomedicals, ref. 6540440) and 500uL of 1x HALT (Thermo Scientific, ref. 78429) in T-PER (Thermo Scientific, ref. 78510). Homogenates were centrifuged at 4 °C and 10,000 xg for 5 minutes. Protein concentration was determined by BCA assay (Thermo Scientific, ref. 23227) using the manufacturer’s protocol. Lysates were normalized and stored at -80 °C until analysis. IGF-1 in liver lysates were quantified by ELISA assay (R&D Systems DuoSet kits) performed according to the manufacturer’s recommendation. Briefly, the capture antibody was coated onto a 96-well half-area plate (Corning) in PBS overnight at room temperature. On the following day, the plates were washed three times with 200 uL of Wash Buffer (0.05% Tween 20 in PBS) and blocked for one hour. After blocking and incubation, plates were washed again before adding samples of small intestinal tissue lysate as well as a standard curve. The plate was incubated for 2 hours at room temperature before being washed as described above. Detection Antibody provided in the kit was added at the suggested concentration and incubated for another 2 hours at room temperature. After incubation, plates were washed and Substrate Solution (1:1 mixture of Color Reagent A (H_2_O_2_) and Color Reagent B (Tetramethylbenzidine)) was added to each well and incubated at room temperature for 20 minutes avoiding direct light. After incubation, Stop Solution (2 N H_2_SO_4_) was added to each well. Plates were reader an optical density of 450 with background subtraction at 570 nm using a Tecan plate reader.

### Statistical Analysis

Statistical analyses were performed in GraphPad Prism (version 10.4.1) unless otherwise noted. V3-V4 16s sequencing data was analyzed in R (version 4.4.2). Statistical details including the number of animals or samples can be found in figure legends. Statistical significance was assessed by Mann-Whitney U test when comparing between two groups, or by Two-Way ANOVA with Šídák’s multiple comparisons test when comparing >2 groups unless otherwise noted. PERMANOVA tests were performed using R package vegan (version 2.6-8) and pairwise comparisons from distance matrices were calculated using the pairwiseAdonis package^44^ (version 0.4.1). P values are shown in figures or tables for samples with significant differences. Each data point represents an individual animal, and horizontal bars represent the mean.

## Supplemental information titles and legends

**Supplemental Figure 1.**
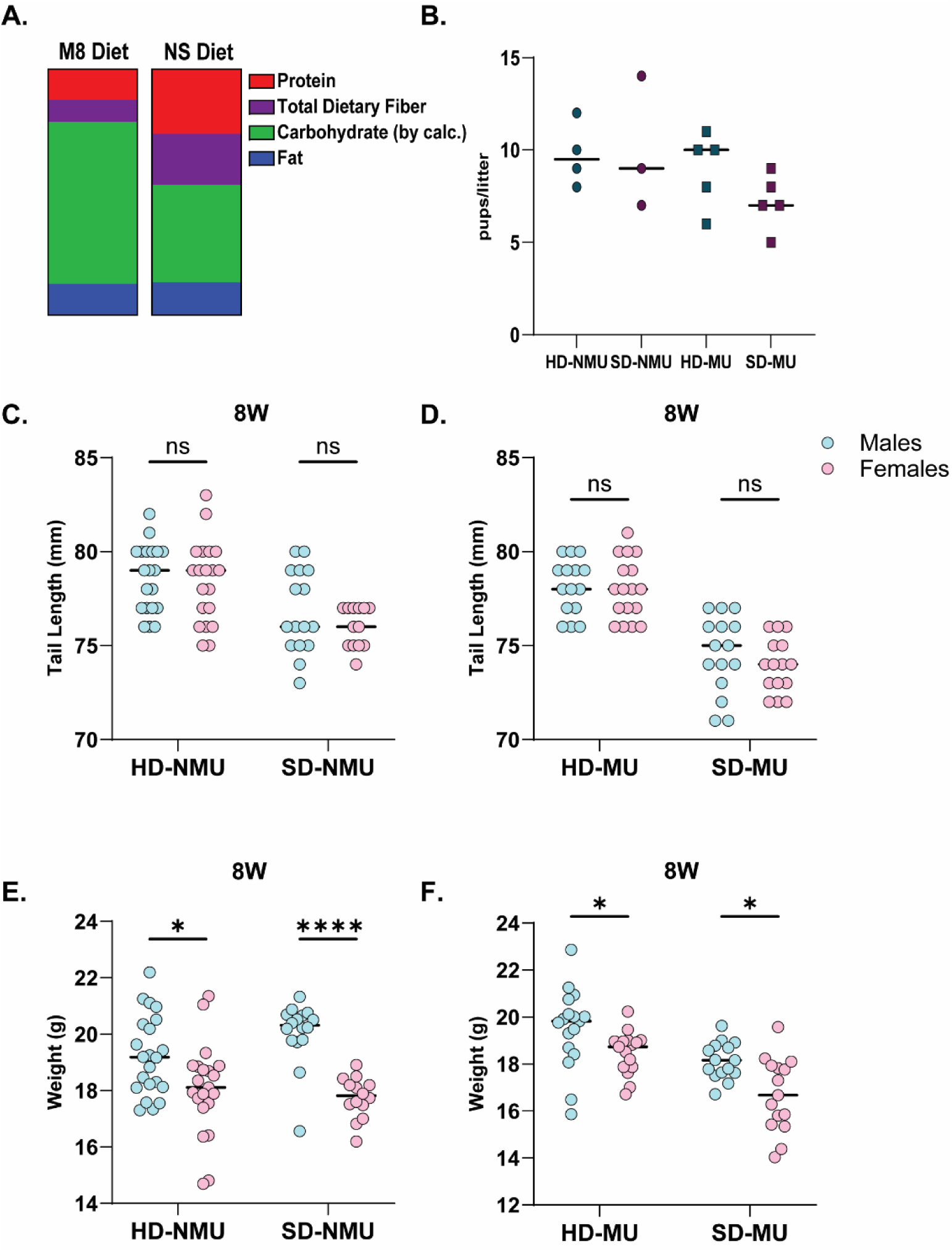
Growth deficits in SD-MU mice are not dependent on litter size or sex. **(A)** Stacked bar plots of major diet components in Malawi-8 (M8) and Nutrient Sufficient diet (NS). **(B)** Number of pups per litter of mice measured for tail length and weight in NMU and MU models at maturity. **(C)** Tail length of NMU mice by sex at maturity. **(D)** Tail length of MU mice by sex at maturity. **(E)** Absolute body weight of NMU mice by sex at maturity. **(F)** Absolute body weight of MU mice by sex at maturity. Each individual point represents a mouse. **(A, C-F)** * p ≤ 0.05, **** p ≤ 0.0001 by Two-Way ANOVA with Holm-Šídák’s multiple comparisons test.

**Figure S2.**
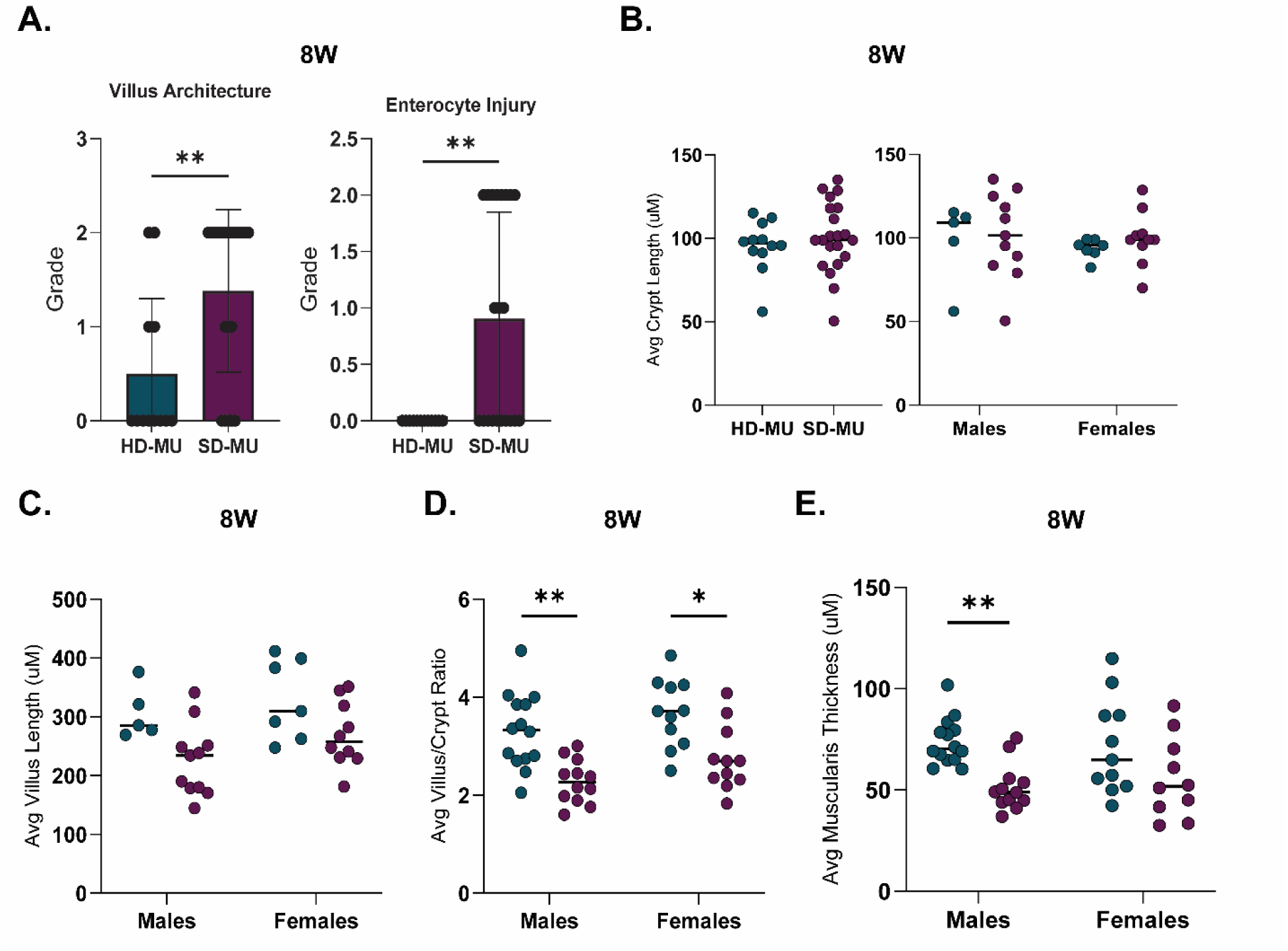
SD-MU offspring exhibit impaired villus architecture and enterocyte injury at maturity. **(A)** Villus Architecture and Enterocyte Injury grade score of H&E-stained ileal tissue from 8-week-old MU animals. **(B)** Average crypt length of H&E-stained ileal tissue from 8-week-old MU animals. Quantification of **(C)** villus length of H&E-stained ileal tissue from 8-week-old MU animals by sex. **(D)** Average villus/crypt ration of H&E-stained ileal tissue from 8-week-old MU animals by sex. **(E)** Average muscularis thickness of H&E-stained ileal tissue from 8-week-old MU animals by sex. Each individual point represents a mouse. * p ≤ 0.05, ** p ≤ 0.01, *** p ≤ 0.001, **** p ≤ 0.0001 by **(A-B)** Mann-Whitney U test or **(B-E)** Two-Way ANOVA with Holm-Šídák’s multiple comparisons test.

**Figure S3.**
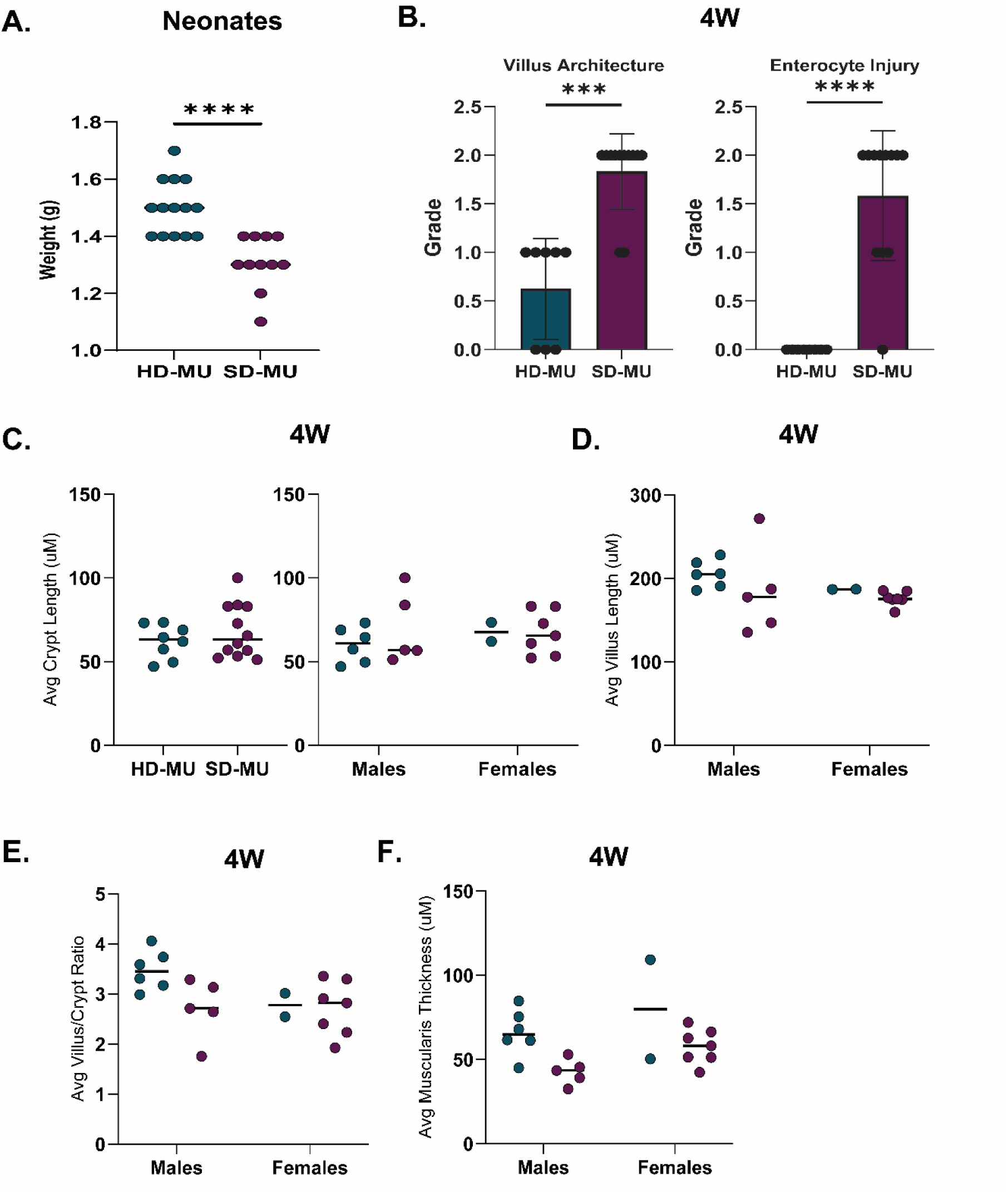
SD-MU offspring exhibit impaired villus architecture and enterocyte injury soon after weaning. **(A)** Absolute body weight of 1-day-old neonates born to HD-MU or SD-MU dams. **(B)** Villus Architecture and Enterocyte Injury grade score of H&E-stained ileal tissue from 4-week-old MU animals. **(C)** Average crypt length of H&E-stained ileal tissue from 4-week-old MU animals. Quantification of **(D)** Villus length of H&E-stained ileal tissue from 4-week-old MU animals by sex. **(E)** Villus/crypt ratio of H&E-stained ileal tissue from 4-week-old MU animals by sex. **(E)** Muscularis thickness of H&E-stained ileal tissue from 4-week-old MU animals by sex. Each individual point represents a mouse. * p ≤ 0.05, ** p ≤ 0.01, *** p ≤ 0.001, **** p ≤ 0.0001 by **(A-C)** Mann-Whitney U test or **(C-F)** Two-Way ANOVA with Holm-Šídák’s multiple comparisons test.

**Figure S4.**
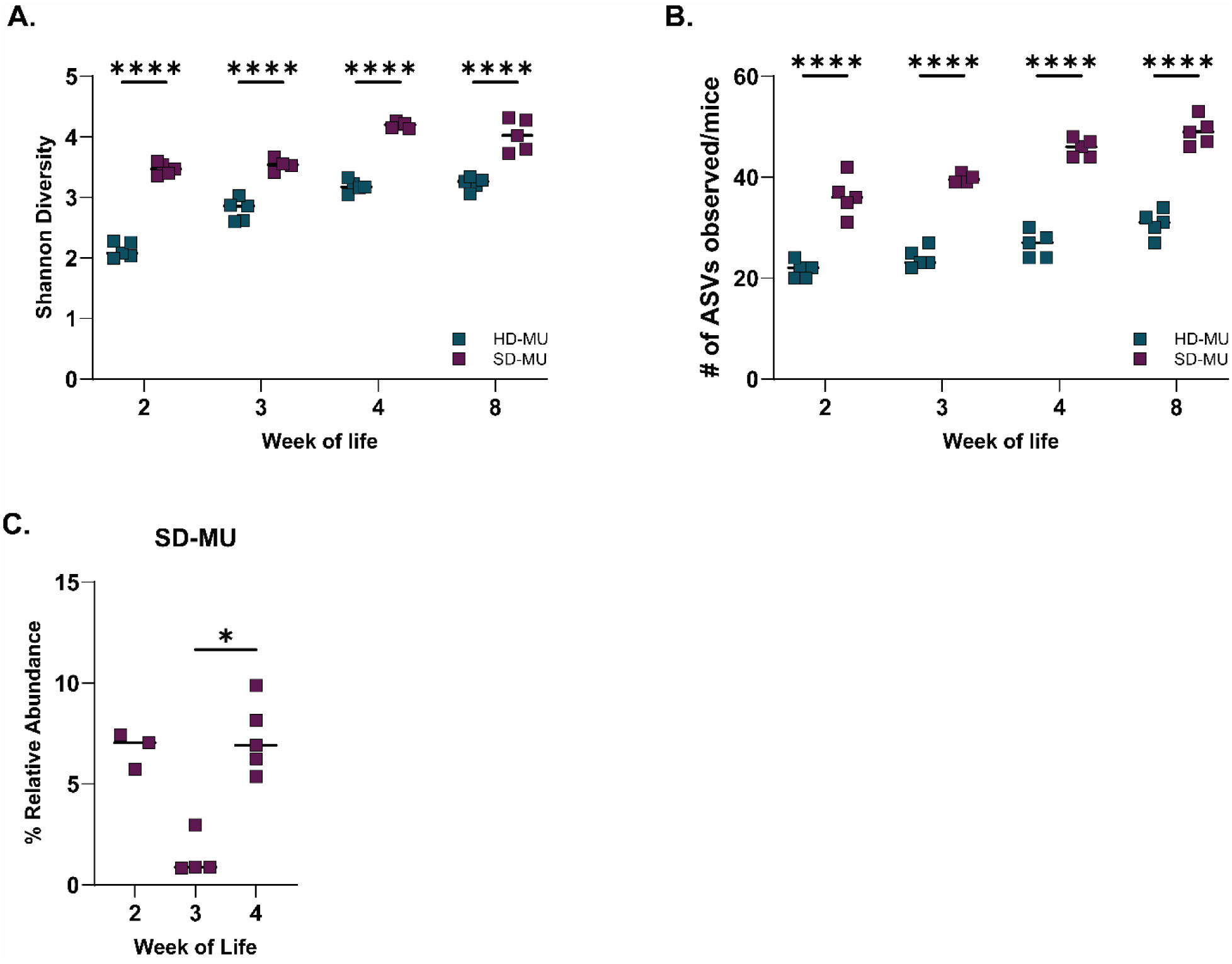
SD-MU Offspring exhibit higher alpha diversity soon after birth. **(A)** Shannon Diversity Index for MU animals in early life. **(B)** Number of ASVs observed in the microbiota of MU animals in early life. **(C)** Percentage relative abundance of *Escherichia coli* before (2-weeks) and after weaning (3-weeks). Each individual point represents a mouse. * p ≤ 0.05, ** p ≤ 0.01, *** p ≤ 0.001, **** p ≤ 0.0001 by **(A-B)** Two-Way ANOVA with Holm-Šídák’s multiple comparisons test or **(C)** Kruskal-Wallis with Dunn’s Test.

**Figure S5.**
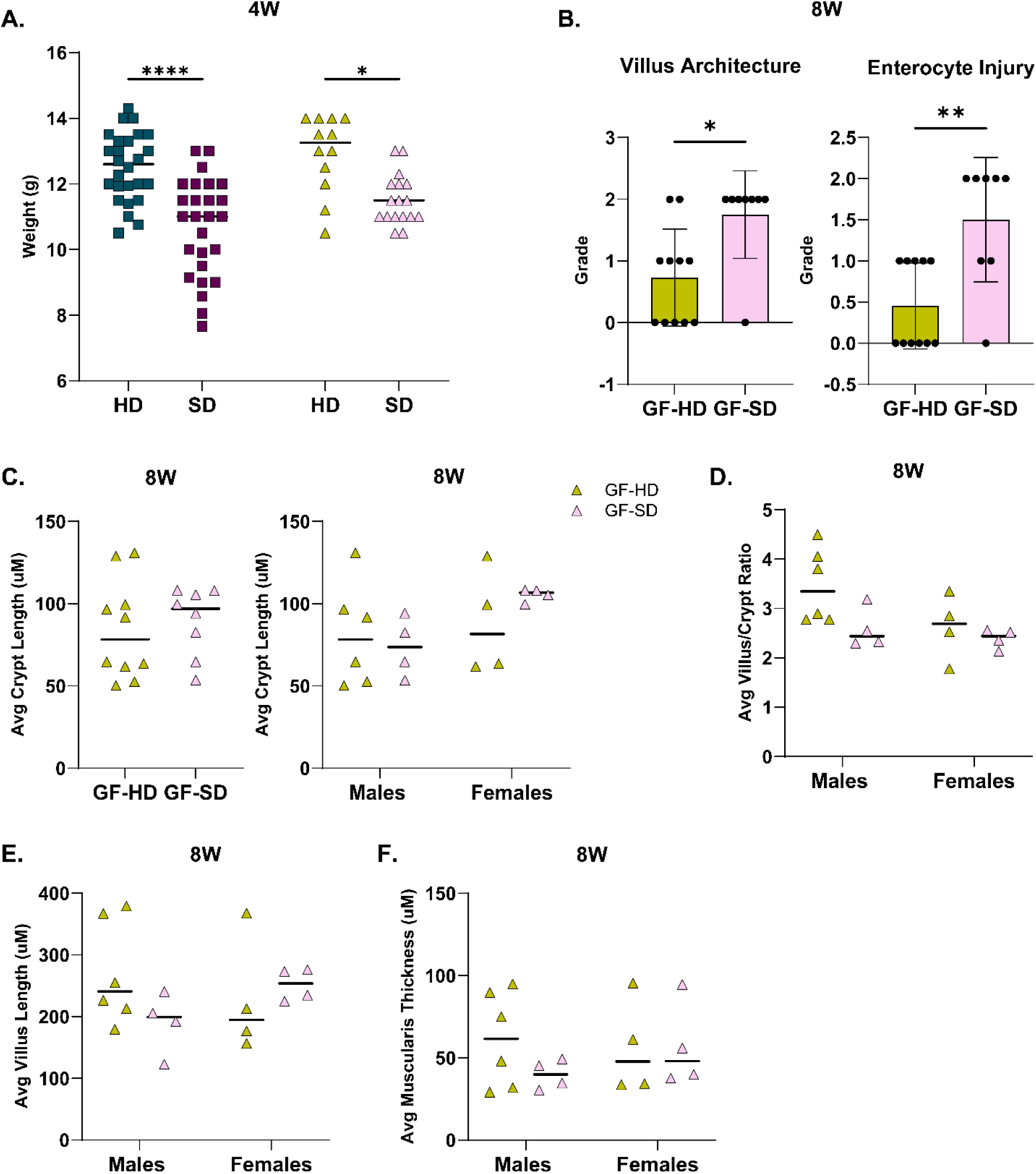
Postnatal factors shape ponderal growth and intestinal morphology. **(A)** Absolute body weight of HD-MU, SD-MU, GF-HD and GF-SD animals 1-week post weaning. **(B)** Villus Architecture and Enterocyte Injury histology score of H&E stained ileal tissue collected from GF-HD and GF-SD at maturity. Quantification of **(C)** crypt length, **(D)** villus/crypt ratio and **(E)** villus length of H&E stained ileal tissue collected from GF-HD and GF-SD at maturity shown by sex. Each point represents an individual mouse. * p ≤ 0.05, ** p ≤ 0.01, *** p ≤ 0.001, **** p ≤ 0.0001 by **(B-C)** Mann-Whitney U test or **(A, C-E)** Two-Way ANOVA with Holm-Šídák’s multiple comparisons test.

**Figure S6.**
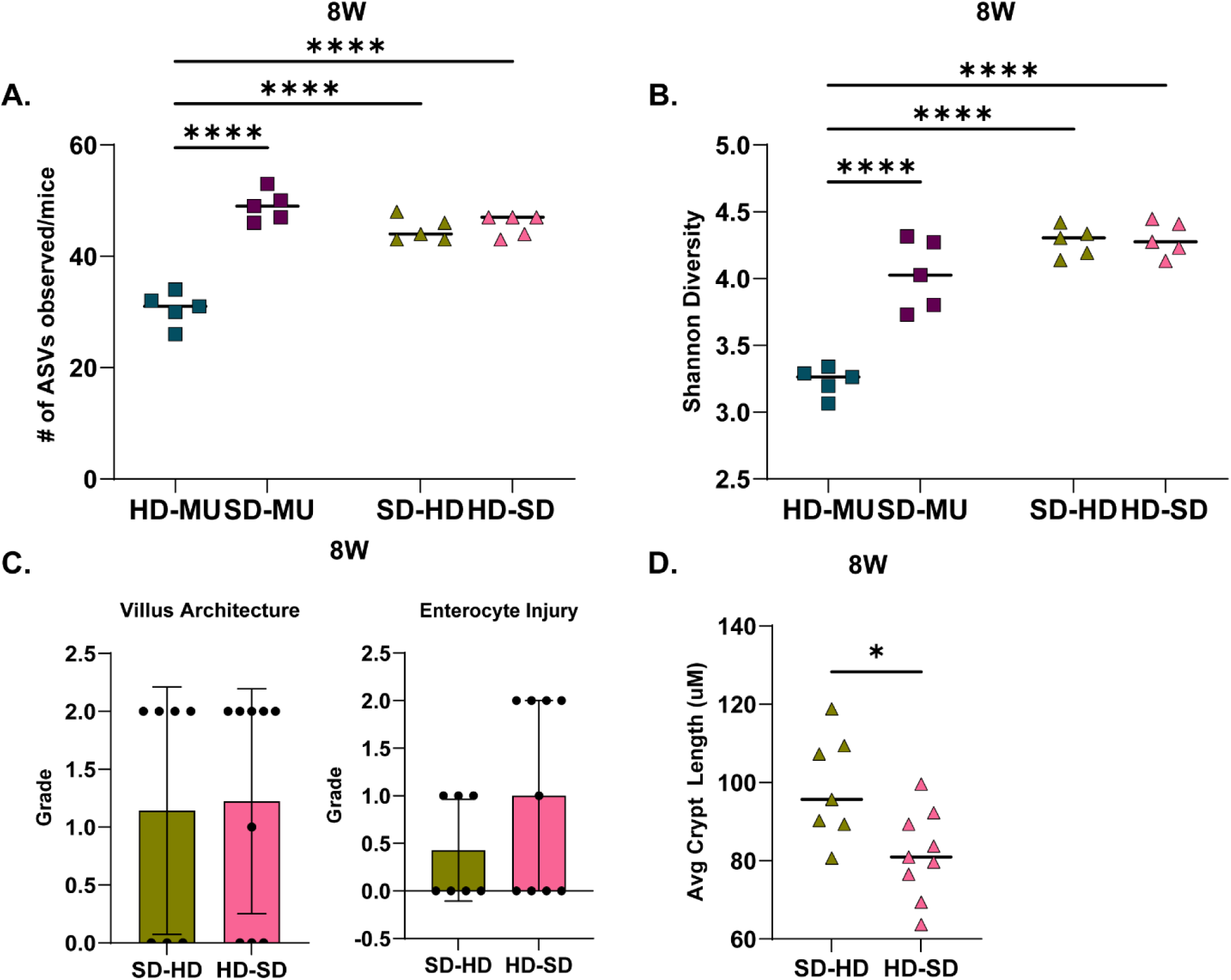
Cross-fostering shapes intestinal microbiota and morphology at maturity. **(A)** Number of ASVs observed in the microbiota of SD-HD and HD-SD animals at maturity. **(B)** Shannon diversity index of cross-fostered and MU animals at maturity. **(C)** Villus Architecture and Enterocyte Injury histology score of H&E stained ileal tissue collected from SD-HD and HD-SD at maturity. **(D)** Quantification of crypt length of H&E stained ileal tissue collected from SD-HD and HD-SD at maturity. Each point represents an individual mouse. * p ≤ 0.05, ** p ≤ 0.01, *** p ≤ 0.001, **** p ≤ 0.0001 by **(A-B)** Two-Way ANOVA with Holm-Šídák’s multiple comparisons test or **(C-D)** Mann-Whitney U test.

## REFERENCES

1. Benjamin-Chung, J. et al. Early-childhood linear growth faltering in low- and middle-income countries. Nature 621, 550–557 (2023).

2. De Sanctis, V. et al. Early and Long-term Consequences of Nutritional Stunting: From Childhood to Adulthood. Acta Biomed. 92, e2021168 (2021).

3. Robertson, R. C., Manges, A. R., Finlay, B. B. & Prendergast, A. J. The Human Microbiome and Child Growth - First 1000 Days and Beyond. Trends Microbiol. 27, 131–147 (2019).

4. Prendergast, A. J. & Humphrey, J. H. The stunting syndrome in developing countries. Paediatr. Int. Child Health 34, 250–265 (2014-4).

5. Cowardin, C. A. et al. Environmental enteric dysfunction: gut and microbiota adaptation in pregnancy and infancy. Nat. Rev. Gastroenterol. Hepatol. (2022) doi:10.1038/s41575-02200714-7.

6. Elisaria, E., Mrema, J., Bogale, T., Segafredo, G. & Festo, C. Effectiveness of integrated nutrition interventions on childhood stunting: a quasi-experimental evaluation design. BMC Nutr 7, 17 (2021).

7. Budge, S., Parker, A. H., Hutchings, P. T. & Garbutt, C. Environmental enteric dysfunction and child stunting. Nutr. Rev. 77, 240–253 (2019).

8. Aziz, I. et al. A prospective study on linking diarrheagenic E. coli with stunted childhood growth in relation to gut microbiome. Sci. Rep. 13, 6802 (2023).

9. Robertson, R. C. et al. The gut microbiome and early-life growth in a population with high prevalence of stunting. Nat. Commun. 14, 1–15 (2023).

10. Subramanian, S. et al. Persistent gut Microbiota immaturity in malnourished Bangladeshi children. Nature 510, 417–421 (2014).

11. Blanton, L. V. et al. Gut bacteria that prevent growth impairments transmitted by microbiota from malnourished children. Science 351, (2016).

12. Smith, M. I. et al. Gut microbiomes of Malawian twin pairs discordant for kwashiorkor. Science 339, 548–554 (2013).

13. Serrano Matos, Y. A., et al. Colonization during a key developmental window reveals microbiota-dependent shifts in growth and immunity during undernutrition. Microbiome 12, 71 (2024).

14. Meyer, A. M. & Caton, J. S. Role of the Small Intestine in Developmental Programming: Impact of Maternal Nutrition on the Dam and Offspring. Adv. Nutr. 7, 169–178 (2016).

15. Kelly, P. et al. Histopathology underlying environmental enteric dysfunction in a cohort study of undernourished children in Bangladesh, Pakistan, and Zambia compared with United States children. Am. J. Clin. Nutr. 120 Suppl 1, S15–S30 (2024).

16. Liu, T.-C., et al. A novel histological index for evaluation of environmental enteric dysfunction identifies geographic-specific features of enteropathy among children with suboptimal growth. PLoS Negl. Trop. Dis. 14, e0007975 (2020).

17. Malique, A. et al. NAD+ precursors and bile acid sequestration treat preclinical refractory environmental enteric dysfunction. Sci. Transl. Med. 16, eabq4145 (2024).

18. Kau, A. L. et al. Functional characterization of IgA-targeted bacterial taxa from undernourished Malawian children that produce diet-dependent enteropathy. Science translational medicine 7, 276ra24–276ra24 (2015).

19. Salameh, E. et al. Modeling undernutrition with enteropathy in mice. Sci. Rep. 10, 15581 (12/2020).

20. Kotloff, K. L. et al. Burden and aetiology of diarrhoeal disease in infants and young children in developing countries (the Global Enteric Multicenter Study, GEMS): a prospective, case-control study. Lancet 382, 209–222 (2013).

21. Sangiorgio, G., Calvo, M., Migliorisi, G., Campanile, F. & Stefani, S. The impact of Enterococcus spp. In the immunocompromised host: A comprehensive review. Pathogens 13, 409 (2024).

22. Nuryandari, S., Widjaja, N. A. & Husada, D. TNF-α and IGF-1 levels in stunting children with chronic infection. Acta Biomed. Ateneo Parmense 95, e2024182 (2024).

23. Ong, K. K. et al. Insulin-like growth factor I concentrations in infancy predict differential gains in body length and adiposity: the Cambridge Baby Growth Study. Am. J. Clin. Nutr. 90, 156–161 (2009).

24. Schwarzer, M. et al. Microbe-mediated intestinal NOD2 stimulation improves linear growth of undernourished infant mice. Science 379, 826–833 (2023).

25. Ahmed, S. et al. Association of Anti-Rotavirus IgA Seroconversion with Growth, Environmental Enteric Dysfunction and Enteropathogens in Rural Pakistani Infants. Vaccine 40, 3444–3451 (2022).

26. Pruss, K. M. et al. Effects of intergenerational transmission of small intestinal bacteria cultured from stunted Bangladeshi children with enteropathy. bioRxivorg 2024.11.01.621574 (2024) doi:10.1101/2024.11.01.621574.

27. Marume, A., Archary, M. & Mahomed, S. Predictors of stunting among children aged 6-59 months, Zimbabwe. Public Health Nutr. 26, 820–833 (2023).

28. Abdulla, F., Rahman, A. & Hossain, M. M. Prevalence and risk predictors of childhood stunting in Bangladesh. PLoS One 18, e0279901 (2023).

29. Lakshmy, R. Metabolic syndrome: role of maternal undernutrition and fetal programming. Rev. Endocr. Metab. Disord. 14, 229–240 (2013).

30. Keats, E. C. et al. Effective interventions to address maternal and child malnutrition: an update of the evidence. *Lancet Child Adolesc*. Health 5, 367–384 (2021).

31. Di Gesù, C. M. et al. Maternal gut microbiota mediate intergenerational effects of high-fat diet on descendant social behavior. Cell Rep. 41, 111461 (2022).

32. Buffington, S. A. et al. Microbial reconstitution reverses maternal diet-induced social and synaptic deficits in offspring. Cell 165, 1762–1775 (2016).

33. Iddrisu, I. et al. Malnutrition and Gut Microbiota in Children. Nutrients 13, 2727 (2021).

34. Rinanda, T., Riani, C., Artarini, A. & Sasongko, L. Correlation between gut microbiota composition, enteric infections and linear growth impairment: a case-control study in childhood stunting in Pidie, Aceh, Indonesia. Gut Pathog. 15, 54 (2023).

35. Dang, H. et al. Maternal gut microbiota influence stem cell function in offspring. Cell Stem Cell 32, 246–262.e8 (2025).

36. Ashorn, P. et al. The impact of lipid-based nutrient supplement provision to pregnant women on newborn size in rural Malawi: a randomized controlled trial. Am. J. Clin. Nutr. 101, 387–397 (2015).

37. Hughes, R. L. et al. Infant gut microbiota characteristics generally do not modify effects of lipid-based nutrient supplementation on growth or inflammation: secondary analysis of a randomized controlled trial in Malawi. Sci. Rep. 10, 14861 (2020).

38. Ashorn, P. et al. Supplementation of maternal diets during pregnancy and for 6 months postpartum and infant diets thereafter with small-quantity lipid-based nutrient supplements does not promote child growth by 18 months of age in rural Malawi: A randomized controlled trial. J. Nutr. 145, 1345–1353 (2015).

39. Cowardin, C. A. et al. Mechanisms by which sialylated milk oligosaccharides impact bone biology in a gnotobiotic mouse model of infant undernutrition. Proc. Natl. Acad. Sci. U. S. A. 116, 11988–11996 (2019).

40. Bodenhofer, U., Bonatesta, E., Horejš-Kainrath, C. & Hochreiter, S. msa: an R package for multiple sequence alignment. Bioinformatics 31, 3997–3999 (2015).

41. Schliep, K. P. phangorn: phylogenetic analysis in R. Bioinformatics 27, 592–593 (2011).

42. McMurdie, P. J. & Holmes, S. phyloseq: an R package for reproducible interactive analysis and graphics of microbiome census data. PLoS One 8, e61217 (2013).

43. Wickham, H. Data Analysis. in Use R! 189–201 (Springer International Publishing, Cham, 2016).

44. Martinez, A. P. PairwiseAdonis: Pairwise Multilevel Comparison Using Adonis. (2020).

